# Integration with a toehold switch expands the dynamic range of a transcription factor-based whole-cell pyruvate biosensor

**DOI:** 10.64898/2026.11.26.769166

**Authors:** Zihan Gao, Maria Suarez-Diez, Pieter Candry

## Abstract

Whole-cell biosensors based on transcription factors (TFs) provide a powerful approach for metabolite sensing, but their dynamic range can be constrained by the coupling of ligand recognition and reporter expression within a single transcriptional regulatory layer. Here, we developed a modular transcription factor–toehold switch (TF–TS) biosensor architecture. This architecture couples TF-mediated metabolite recognition to downstream translational activation with a toehold switch as an additional RNA regulatory layer. Using two rounds of Design–Build– Test–Learn cycles guided by Design of Experiments, we systematically optimized the toehold switch module by tuning plasmid copy number, ribosome binding site (RBS) strength, and inducer-controlled RNA expression. A medium-copy-number construct with a strong RBS achieved the highest ON/OFF ratio of 33.70 ± 2.36. This optimized module was coupled to the pyruvate-responsive transcription factor PdhR. The resulting Pyr-toehold biosensor showed a 44.7-fold higher dynamic range and a 210-fold lower basal expression compared with the conventional PdhR-based biosensor. Temporal induction experiments further demonstrated that the two regulatory layers could be activated independently, enabling modulation of the timing and magnitude of reporter activation. Together, these results demonstrate that coupling TF-mediated metabolite recognition with programmable RNA-level translational control can improve both the dynamic range and temporal controllability of whole-cell biosensors. This modular architecture provides a potential framework for engineering biosensors with independently tunable signal output and regulatory dynamics.

## 1. Introduction

Living bacterial cells innately sense and respond to environmental changes(*1*). Synthetic biology takes advantage of this capability by engineering genetic circuits that convert chemical or physical signals into measurable outputs, such as fluorescence(*2*). Such bacterial whole-cell biosensors offer rapid responses and are more cost-effective and environmentally friendly than conventional methods. This has led to diverse applications, such as metabolic engineering(*3, 4*), environmental monitoring(*5*), food safety detection(*6*) and disease diagnosis(*7*). Among these synthetic biosensors, transcription factor (TF)-based biosensors are among the most extensively developed and widely adopted platforms(*8, 9*). Ligand-responsive TFs are proteins that can bind a ligand as well as DNA; during ligand-binding, TF undergo conformational changes that affects DNA binding, in turn activating or repressing downstream gene expression(*10*). Their wide adoption stems from the abundance of naturally occurring ligand-responsive TFs, their high sensitivity and specificity, straightforward genetic implementation, and seamless integration with endogenous transcriptional regulatory networks(*11–13*).

For applications such as precise metabolic regulation and quantitative sensing, TF-based biosensors require a high dynamic range to function effectively. A high dynamic range improves the signal-to-background ratio, enables the discrimination of subtle changes in ligand concentration, and facilitates the accurate identification of high-producing strains during high-throughput screening(*14*). In general, biosensor dynamic range can be improved by reducing basal expression and/or increasing maximal output(*15*). To this end, numerous engineering strategies have been developed, including promoter engineering(*16, 17*), ribosome binding site (RBS) optimization(*18, 19*), TF engineering(*20*), genetic amplifier circuits(*21*), and plasmid copy number (CN) optimization(*22*). Promoter engineering can target either the promoter driving TF expression, or the TF-responsive output promoter. Engineering of the latter promoter regulates transcription initiation by modulating both RNA polymerase recruitment and TF binding to operator sequences, thereby influencing both basal leakage and maximal output (*23, 24*). Similarly, RBS strength modulates ribosome recruitment and translation efficiency, making it a key determinant of the translation initiation rate of output mRNAs. Consequently, RBS engineering directly affects leakiness and maximal output of the biosensor (*25, 26*). Although these approaches can improve biosensor performance, they generally modify individual regulatory components without changing the underlying signal-transduction architecture, retaining the coupling between basal and inducible expression (*27*).

To address these limitations, we introduced an additional layer of translational regulation by coupling a TF-based biosensor with a programmable toehold switch. This approach enables regulation at both the transcriptional and translational levels, providing an additional mechanism to suppress basal expression and thereby expand the dynamic range. Toehold switches are synthetic RNA-based translational regulators that control gene expression through trigger RNA-induced rearrangement of RNA secondary structure (*28, 29*). Upon hybridization with a sequence-complementary trigger RNA, the hairpin structure unfolds, exposing the RBS and initiating translation(*30*). Their programmability and orthogonality have enabled applications in RNA sensing and pathogen diagnostics, such as Zika virus and SARS-CoV-2 detection(*31, 32*). However, conventional toehold switches respond to RNA triggers and cannot directly recognize small molecules. We hypothesize that integrating a TF-controlled promoter with a toehold switch provides a route to convert metabolite recognition into an RNA signal, thereby extending toehold-switch-based sensing to small molecules. Combining these transcriptional and translational regulatory layers could reduce basal expression and increase the dynamic range of the biosensor.

In this work, we first established a tunable dual-input toehold switch–trigger (TSTr) system and applied a Design–Build–Test–Learn (DBTL)-guided engineering framework integrating Design of Experiments (DoE) to systematically optimize its performance. As rapid reporter turnover is important for monitoring dynamic metabolic processes(*33, 34*), a second DBTL cycle was used to investigate the effect of the degradation tag fused to the superfolder green fluorescent protein (sfGFP) reporter on signal decay. The optimized toehold switch module was integrated with ligand-responsive transcription factors to construct a transcription factor-toehold switch (TF– TS) hybrid biosensor. Using pyruvate as a model metabolite, which serves as a central intermediate linking glycolysis and the tricarboxylic acid (TCA) cycle and as a key precursor for the biosynthesis of valuable chemicals(*35, 36*), we coupled the toehold switch module with the pyruvate-responsive system, in which pyruvate relieves PdhR-mediated repression of P_pdh_(*37, 38*). The resulting TF–TS hybrid pyruvate biosensor exhibited an expanded dynamic range, reduced basal expression, and tunable temporal response. More broadly, this work establishes a modular and programmable toehold switch platform that can be integrated with TF-based biosensors to coordinate transcriptional and translational regulation, improving the dynamic range and temporal response of whole-cell biosensors.

## 2. Materials & methods

### 2.1 Strains, media and cultivation conditions

All bacterial strains used in this study are listed in Supplementary Table S1. *Escherichia coli* DH5α was used for cloning, DNA assembly and plasmid construction. *E. coli* BL21(DE3) was used for preliminary validation of the toehold switch system and RBS function. *E. coli* Marionette MG1655 was used for TSTr-series constructs and hybrid biosensor performance characterization(*39*).

For characterization experiments, individual colonies were grown overnight in Lysogeny broth (LB) medium supplemented with ampicillin or apramycin as appropriate at 37 °C and 250 rpm. Cultures were then washed twice with M9 minimal medium and adapted for 4 h in M9 medium containing 10 mM glucose and the corresponding antibiotic. Cells were subsequently washed once and diluted to an initial OD_600_ of 0.1 before use. Detailed medium compositions and stock solution preparation are provided in Supplementary Method S1.

### 2.2 Biosensor genetic circuit construction and transformation

All plasmids, genetic parts, sequences, and primers used in this study are listed in Tables S2–S4, and plasmid maps are provided in Figure S1. pUC19 (ColE1, ampicillin resistance) and pSEVAb86 (p15A, apramycin resistance) were used as high- and medium-copy-number vectors, respectively, with reported copy numbers of ∼61 and ∼9 per cell(*40*). Toehold switch and trigger RNA sequences were obtained from a previous study(*29*), and constitutive promoters and RBSs were obtained from the iGEM Parts Registry (https://parts.igem.org/Promoters).

DNA oligonucleotides and promoter/RBS fragments were synthesized by Integrated DNA Technologies (IDT) and assembled into the corresponding vectors using HiFi DNA Assembly (NEB, #E2621). PCR products were treated with DpnI (NEB, #R0176S), and constructs were verified by colony PCR and Sanger sequencing. Plasmids were introduced into the host strains by electroporation using a Gene Pulser Xcell electroporator (Bio-Rad; 2.5 kV, 0.1 cm cuvette). Transformants were selected on LB agar containing the appropriate antibiotics, screened by colony PCR, and further validated by whole-plasmid sequencing (Plasmidsaurus). Confirmed strains were stored as glycerol stocks at −80 °C.

### 2.3 Evaluation of the dual-input TSTr system

In the dual-input TSTr system, P_tet_ and P_cymRC_ controlled expression of the toehold switch and trigger RNAs, respectively, through aTc-inducible TetR and cumate-inducible CymR^AM^ regulation in *E. coli* Marionette MG1655. Cultures were grown in M9 medium containing 10 mM glucose and apramycin. For initial characterization, cells were subjected to four induction conditions: no inducer, aTc alone, cumate alone, or both inducers combined. For pre-culture experiments, cells were pre-cultured for 4 h with either inducer and washed twice before subsequent induction. Dose–response measurements were performed using different concentrations of aTc and cumate.

Each well of a 96-well plate contained 150 µL of culture, overlaid with 50 µL of mineral oil (Sigma-Aldrich, #M8410) to prevent evaporation. Well plates were incubated at 37 °C with shaking at 731 cpm (2 mm orbit). Fluorescence intensity of sfGFP (excitation, 485 nm; emission, 528 nm; fluorescence gain, 65) and optical density at 600 nm (OD_600_) were measured using a microplate reader (BioTek Synergy H1). Biomass growth was accounted for by normalizing GFP fluorescence to OD_600_. Background fluorescence was corrected by subtracting the GFP/OD value of the negative host-strain control. Background-corrected GFP/OD values below zero were set to zero. Cultures were monitored for 12 h, and measurements at 5 h were used for analysis.

### 2.4 DoE-guided optimization of the TSTr-series constructs

A 2^2^ full factorial design was used to evaluate the effects of RBS strength and CN on TSTr-series constructs performance. Strong RBS and high-copy-number ColE1 were each coded as +1, whereas RBS33 and medium-copy-number p15A were each coded as −1 (Table S5). TSTr-series constructs were cultured in M9 medium containing 10 mM glucose and the appropriate antibiotics. The OFF state was induced with 0.2 μM aTc alone, whereas the ON state was induced with 0.2 μM aTc and 100 μM cumate. sfGFP fused to an ASV degradation tag (half-life approx. 110 min (*41*)) was used as the fluorescent reporter. Cultures were incubated and fluorescence was measured under the conditions described above.

A 2^3^ full factorial design was used to evaluate the effects of CN (p15A, −1; ColE1, +1), aTc concentration (0.01, −1; 0.02 μM, +1), and cumate concentration (10, −1; 100 μM, +1) on signal decay, while the RBS was fixed at strong level. GFP fluorescence was monitored over time, and measurements collected after the maximum fluorescence were fitted to a one-phase decay model.

### 2.5 Characterization of the TF–TS hybrid pyruvate biosensor

The TF–TS hybrid pyruvate biosensor was constructed using the native PdhR-regulated P_pdh_ promoter (sequence provided in Table S3), in which pyruvate relieves PdhR-mediated transcriptional repression. For the Pyr-trigger construct, aTc was added at a final concentration of 0.2 μM to induce toehold switch RNA expression, whereas the Pyr-toehold construct was induced with 100 μM cumate to activate the trigger RNA expression. Pyruvate was added to final concentrations of 0, 0.01, 0.1, 1, 2, 5, 10, and 100 mM. Cultures were incubated and fluorescence was measured under the same conditions described above.

### 2.6 Temporal response analysis of the Pyr-toehold biosensor

Temporal responses of the Pyr-toehold biosensor were monitored using a BioTek Synergy H1 microplate reader equipped with an automated reagent injector for timed addition of pyruvate or cumate. Three induction schemes were tested: (i) cells cultured with 10 mM pyruvate, followed by cumate addition (100 μM) at 0, 2, 4, or 6 h; (ii) cells cultured with 100 μM cumate, followed by pyruvate addition (10 mM) at 0, 2, 4, or 6 h; and (iii) cells cultured with 10 mM glucose, followed by cumate addition (100 μM) at 0, 2, 4, or 6 h. The corresponding no-cumate or no-pyruvate condition was included as a control for each experiment. For each experiment, 135 μL of culture (initial OD_600_ = 0.1) was loaded into each well, followed by injection of 15 μL inducer solution at a dispensing rate of 250 µL/sec. Wells were overlaid with 50 μL mineral oil, and GFP fluorescence and OD_600_ were monitored for 12 h under the conditions described above.

### 2.7 Data processing and statistical analysis

All measurements were performed using three biologically independent replicates. A two-tailed, unpaired Student’s t-test assuming equal variances was performed using Microsoft Excel 16.80. A *P value* < 0.05 was considered statistically significant. DoE data were analyzed in R 4.3.2 using linear models, and regression coefficients, interaction effects, and adjusted R² were used to evaluate factor effects. Pareto plots were generated using the paretoPlot function(*42*). The ON/OFF ratio was calculated as the ratio between the fluorescence signal under ON-state and OFF-state conditions. For the TSTr-series constructs, the ON state was defined as co-expression of the toehold switch and trigger RNAs, whereas the OFF state was defined as expression of the toehold switch RNA in the absence of trigger RNA. GFP signal decay was fitted using a one-phase decay model to determine the decay rate constant (*K*) and decay fraction (Eq. 1):

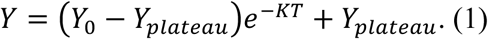

where *T* represents time, *Y* represents the GFP fluorescence signal, *T_0_* represents the time at which the maximum GFP fluorescence signal is reached, *Y*_0_ represents the GFP fluorescence signal at *T* = *T_0_*, *Y_plateau_* represents the asymptotic GFP signal, and *K* represents the decay rate constant. The decay fraction was calculated as (Eq. 2):

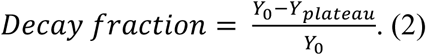

Dose–response data were fitted to the Hill equation (Eq. 3):

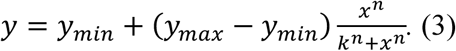

Where *y* is the relative fluorescence intensity (GFP/OD), *y_max_*/*y_min_* is the maximum/minimum relative fluorescence intensity, *x* is the concentration of pyruvate, *k* is the threshold of the gene circuit and *n* is the Hill coefficient. The dynamic range was calculated as the ratio of the fitted maximum and minimum fluorescence values.

Parameter estimation and curve fitting for the Hill (with 1/Y^2^ weighting) and one-phase decay models were performed in GraphPad Prism 10. Detailed equations and model descriptions for the DoE analysis are provided in Method S2. The schematic illustrations in Figures 1A, 2A, 3A and 4A were created with BioRender (BioRender.com).

**Figure 1.**
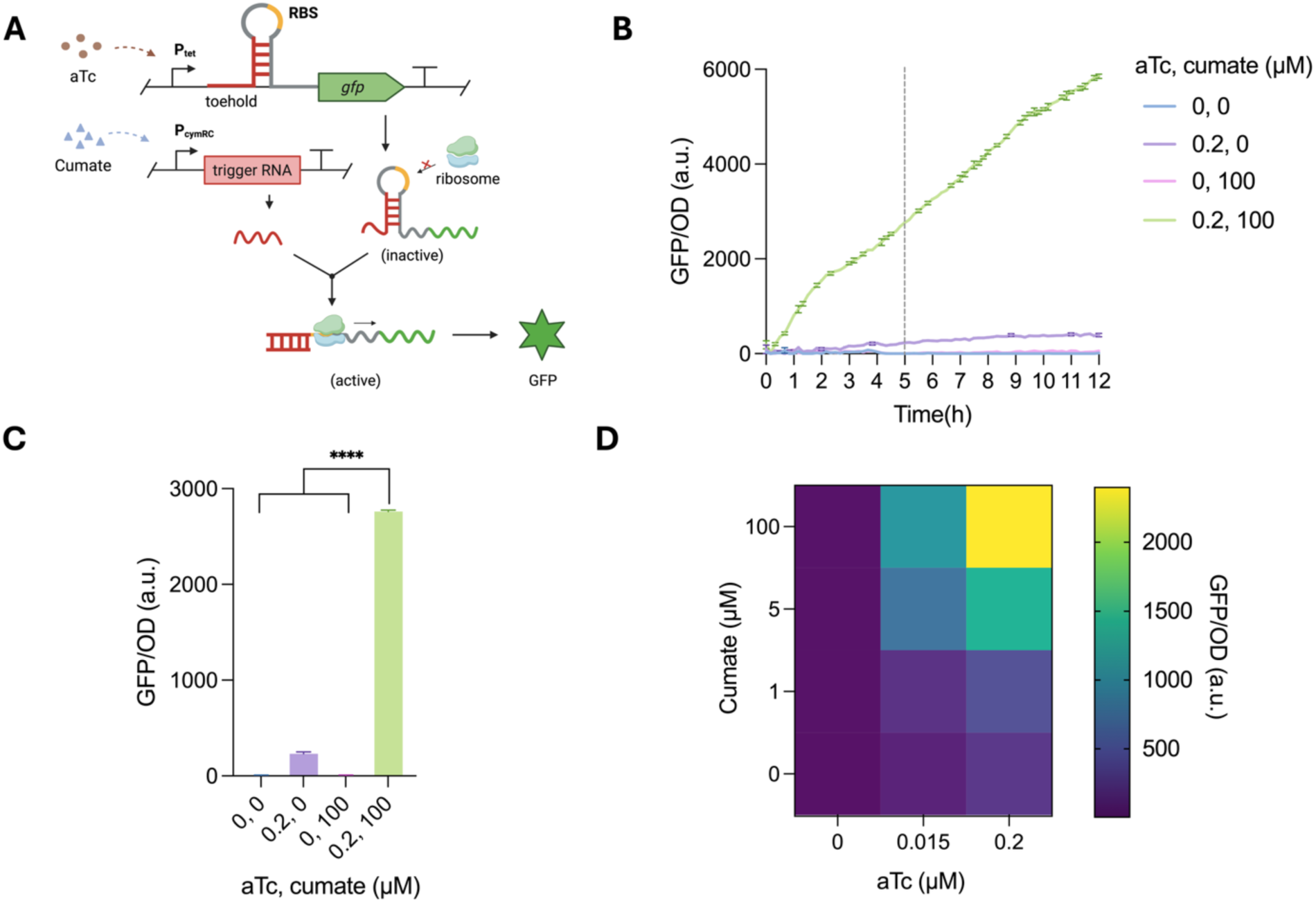
Characterization of AND-gate behavior and tunability of the dual-input TSTr system. (A) Schematic illustration of the dual-input toehold switch system. The expression of toehold switch RNA and trigger RNA was independently regulated by the inducible promoters P_tet_ and P_cymRC_ in response to aTc and cumate, respectively. The *gfp* and GFP labels in all schematic illustrations refer to the sfGFP reporter. (B) Time-course GFP fluorescence under different inducer combinations. (C) GFP fluorescence at 5 h under different inducer combinations, ****P < 0.0001. (D) Heatmap analysis of GFP output under different concentrations of aTc and cumate at 5 h. Data are presented as mean ± SD from three independent biological replicates (n = 3).

**Figure 2.**
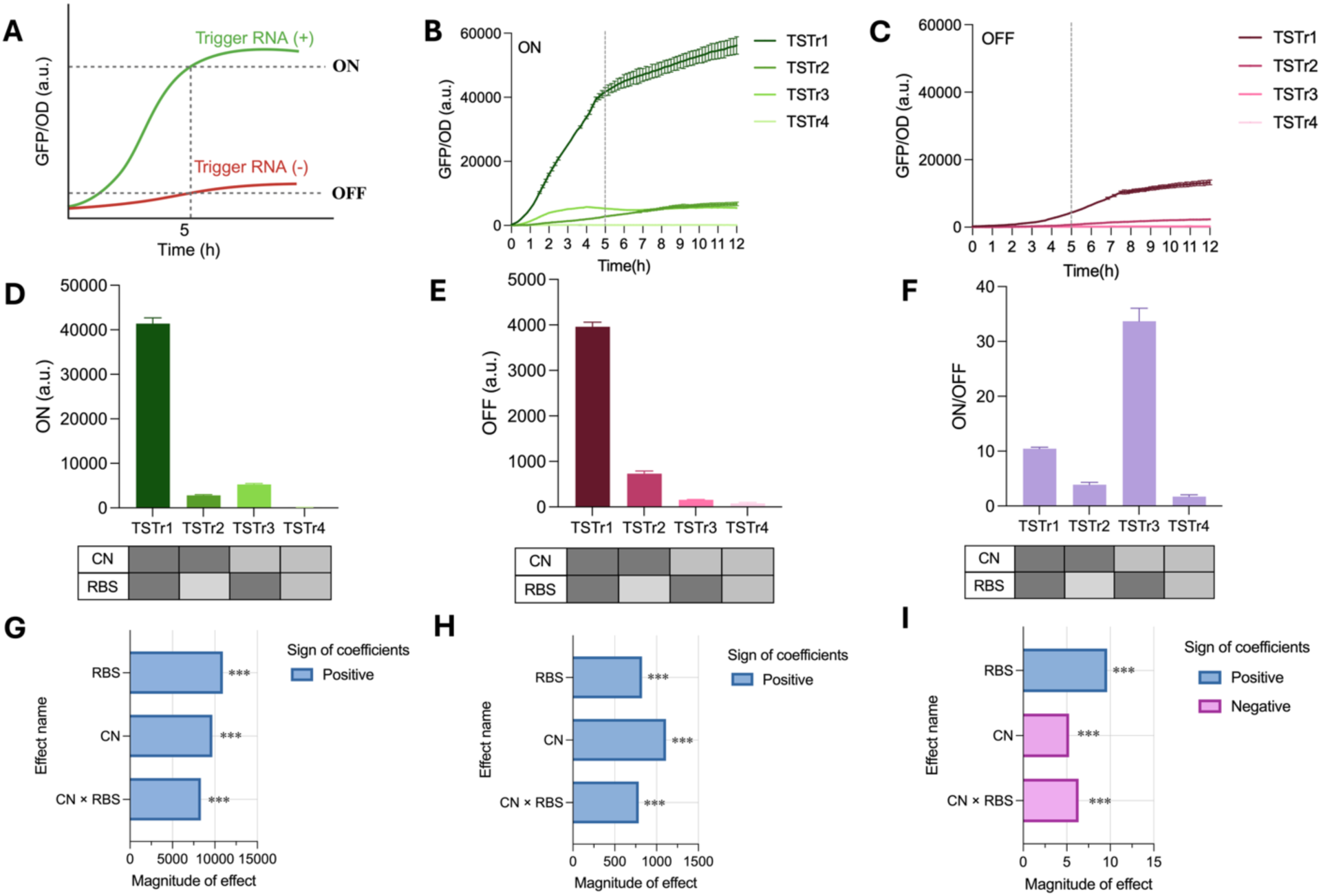
DoE-guided optimization of TSTr-series constructs by tuning plasmid copy number and RBS strength. (A) Schematic illustration of the ON and OFF states used for factorial analysis. (B) Time-course GFP fluorescence under ON-state conditions for the four constructs. (C) Time-course GFP fluorescence under OFF-state conditions. (D) ON-state GFP fluorescence at 5 h. (E) OFF-state GFP fluorescence at 5 h. (F) ON/OFF ratios of the four constructs. The grayscale matrices below panels D–F indicate the plasmid CN and RBS strength of each construct, with dark gray representing the high (+1) level and light gray representing the low (−1) level. (G–I) Regression coefficients for RBS strength, CN, and their two-factor interaction (CN × RBS) on ON-state fluorescence (G), OFF-state fluorescence (H), and the ON/OFF ratio (I).

**Figure 3.**
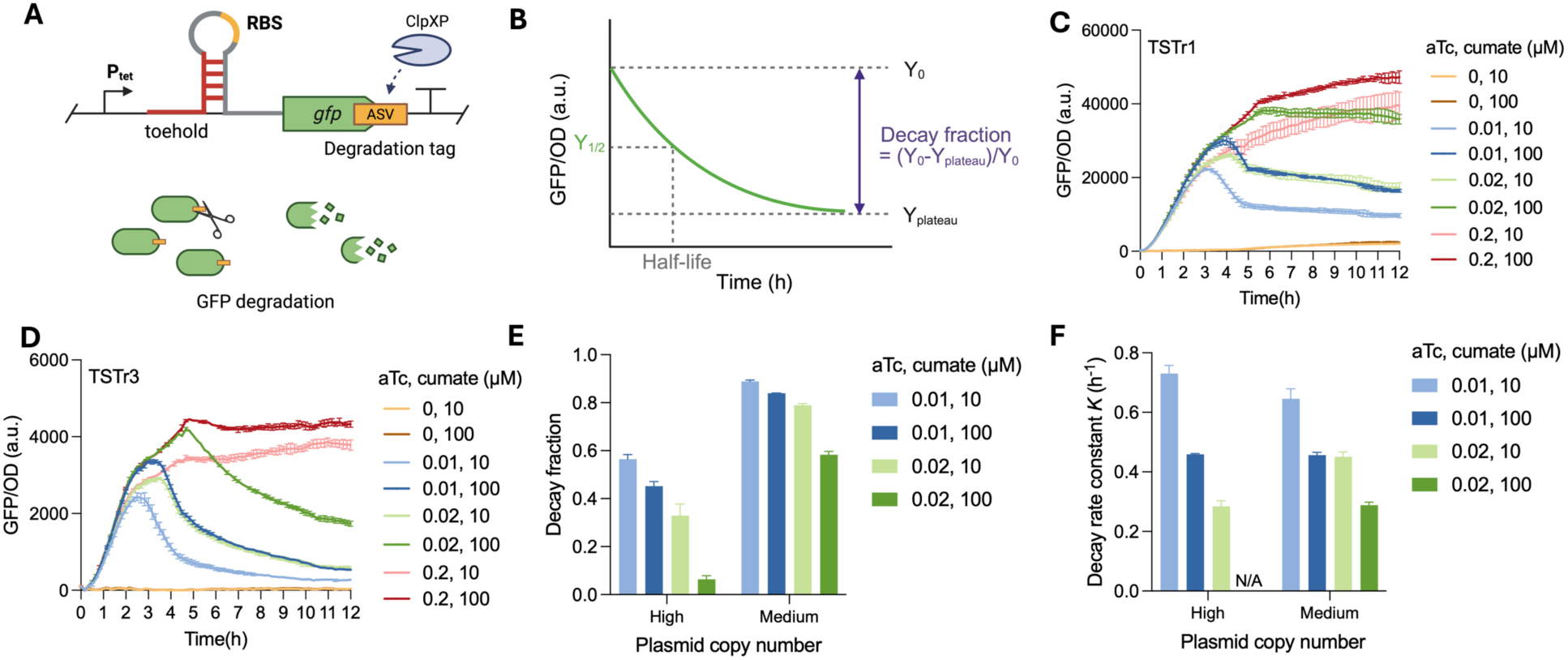
Characterization and optimization of signal decay. (A) Schematic illustration of ASV-tagged *sfGFP* degradation by ClpXP. (B) Schematic illustration of the one-phase decay model. (C–D) Time-course GFP fluorescence of TSTr1 (C) and TSTr3 (D) under different combinations of aTc and cumate concentrations. (E–F) Signal decay fraction (E) and decay rate constant *K* (F) under different combinations of plasmid copy number, aTc concentration, and cumate concentration.

**Figure 4.**
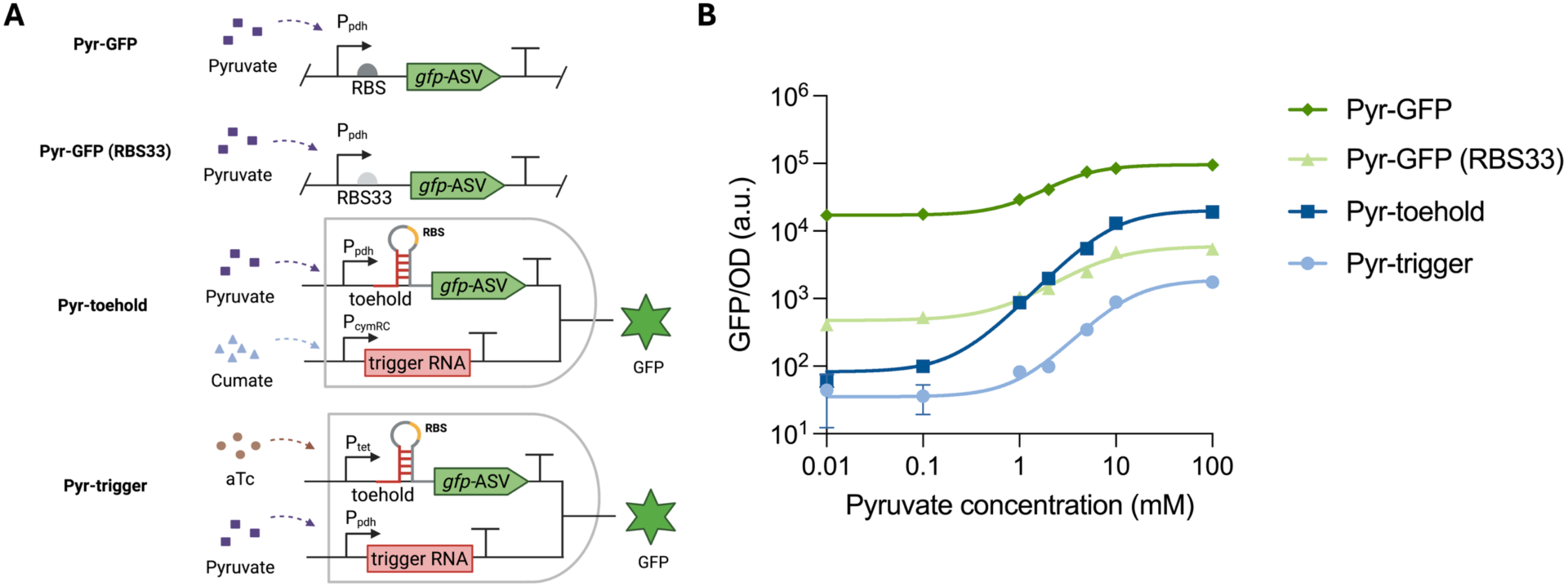
TF-based and TF–TS hybrid architectures for pyruvate biosensing. (A) Design of TF-based and TF–TS hybrid pyruvate biosensor architectures. (B) Pyruvate dose–response curves of the four constructs at 5 h. Solid lines represent Hill equation fits.

## 3. Results

### 3.1 Dual-input TSTr system exhibits tunable AND-gate behavior

The functionality of the toehold switch system was initially validated in *E. coli* BL21(DE3), where both strong- and weak-RBS constructs exhibited low basal expression and robust trigger-dependent activation. Modulating RBS strength primarily affected the ON-state output, while NUPACK predictions supported the expected structural response upon trigger binding (Figures S2, S3). To investigate tunable AND-gate behavior, we used *E. coli* Marionette MG1655 to independently control toehold switch and trigger RNA expression through the aTc-inducible P_tet_ and cumate-inducible P_cymRC_ promoters, respectively (Figure 1A). Based on the dose–response profiles (Figure S4), 0.2 μM aTc and 100 μM cumate were selected as saturating concentrations. In the absence of both inducers, GFP fluorescence remained at a low level, and addition of either aTc or cumate alone resulted in only weak fluorescence. In contrast, simultaneous induction with both aTc and cumate resulted in strong GFP activation, indicating that the system exhibited AND-gate behavior through the independent regulation of both RNA components by two chemical inducers (Figure 1B). At 5 h, GFP fluorescence under dual-inducer conditions was significantly higher than that under either single-inducer or non-induced conditions (Figure 1C), demonstrating AND-gate behavior of the dual-input TSTr system. Pre-culture experiments further demonstrated an asymmetric response: aTc pre-induction enabled subsequent activation by cumate alone, whereas cumate pre-induction did not result in appreciable activation following subsequent aTc addition (Figure S5). Together with the dual-inducer response, these results further support the modular AND-gate behavior of the TSTr system.

The tunability of the biosensor was assessed by varying the concentrations of aTc and cumate. The concentration of aTc was set to 0, 0.015, and 0.2 μM, with 0.015 μM selected close to the EC_50_ of the P_tet_ system (0.0132 μM; Figure S4). Similarly, cumate concentrations were set to 0, 1, 5, and 100 μM, with 5 μM selected close to the EC_50_ of the P_cymRC_ system (4.33 μM; Figure S4). In the absence of aTc, GFP fluorescence remained at a low level across all cumate concentrations. Increasing aTc from 0 to 0.015 μM enabled biosensor activation, after which increasing cumate concentrations progressively increased GFP output. When aTc was further increased from 0.015 to 0.2 μM, the effect of cumate became more pronounced, with higher cumate levels resulting in progressively stronger GFP output. The highest fluorescence output was observed at 0.2 μM aTc combined with 100 μM cumate (Figure 1D). Together, these results demonstrate that the dual-input TSTr system exhibits tunable AND-gate behavior, with aTc primarily controlling system activation and cumate modulating the magnitude of the output after activation.

### 3.2 A strong RBS and medium plasmid copy number maximize the ON/OFF ratio

Following establishment of the dual-input TSTr system, we initiated the first DBTL cycle to optimize its ON/OFF performance by exploring plasmid CN and RBS strength. A 2^2^ full factorial design comprising two CN levels (medium, −1; high, +1) and two RBS strengths (weak, −1; strong, +1) generated four constructs (TSTr1–4; Table S5). Based on the finding that aTc primarily controlled system activation whereas cumate modulated output magnitude, the ON state was induced with 0.2 μM aTc and 100 μM cumate, while the OFF state was induced with 0.2 μM aTc alone, allowing trigger-dependent activation to be assessed while maintaining toehold switch RNA induction (Figure 2A).

The four constructs resulted in markedly different ON-state and OFF-state levels (Figure 2B–E). For the ON state, TSTr1 (high CN, strong RBS) exhibited the highest fluorescence, whereas TSTr4 (medium CN, weak RBS) showed minimal GFP fluorescence, with TSTr2 (high CN, weak RBS) and TSTr3 (medium CN, strong RBS) displaying intermediate levels (Figure 2B, 2D). A similar trend was observed in the OFF state (Figure 2C, 2E). Regression analysis showed significant positive effects of RBS, CN and their two-factor interaction (CN × RBS) on ON- and OFF-state fluorescence, although the magnitudes of these effects differed between the two states (Figure 2G, 2H; Table S6). RBS strength had the largest effect on ON-state fluorescence, whereas CN had the largest effect on OFF-state fluorescence. This suggests that the use of a high-copy- number plasmid contributes strongly to increased basal expression. Overall, increasing either CN or RBS strength increased both induced GFP expression and basal expression in the absence of trigger RNA.

The ON/OFF ratio provides a measure of the separation between trigger-activated and basal output, revealing trade-offs between maximal output and basal expression. Despite producing the highest ON-state fluorescence, TSTr1 did not exhibit the highest ON/OFF ratio because its high plasmid copy number also resulted in substantial OFF-state fluorescence (Figure 2F). TSTr3 achieved the highest ON/OFF ratio of 33.70 ± 2.36, approximately 3.2- and 19.8-fold higher than TSTr1 (10.46 ± 0.27) and TSTr4 (1.71 ± 0.35), respectively (Figure 2F, Table S5). Factorial analysis revealed a positive effect of RBS strength, but negative effects of plasmid copy number and the CN × RBS interaction on the ON/OFF ratio (Figure 2I; Table S6). These results indicate that the strong RBS enhanced inducible output, whereas increasing CN reduced the ON/OFF ratio by increasing basal expression. These findings identified TSTr3 as the construct with the highest ON/OFF ratio in the first DBTL cycle. Based on the positive effect of RBS strength, TSTr1 and TSTr3, both containing the strong RBS but differing in CN, were selected for the next optimization cycle.

### 3.3 Plasmid copy number and inducer concentration influence signal decay

To test the temporal response of the TSTr system, we evaluated signal decay using the previously established sfGFP reporter fused to an ASV degradation tag, which promotes ClpXP-mediated reporter degradation in *E. coli*(*43*) (Figure 3A). Signal decay was characterized using a one-phase decay model, from which the decay fraction and decay rate constant (*K*) were determined (Figure 3B). To improve suitability for dynamic sensing applications, the second DBTL cycle investigated how CN and inducer concentrations influenced signal decay in TSTr1 and TSTr3.

We first evaluated aTc and cumate concentrations to identify induction conditions that resulted in measurable signal decay. aTc concentrations of 0, 0.01, 0.02, and 0.2 μM were used to modulate toehold switch RNA expression, while cumate concentrations of 10 and 100 μM were used to vary trigger RNA expression (Figure 3C, 3D). At 0 μM aTc, almost no detectable GFP fluorescence was observed for either construct. At intermediate aTc concentrations (0.01 and 0.02 μM), induction with 10 or 100 μM cumate generated pronounced GFP signals followed by measurable signal decay, although the extent of decay varied between TSTr1 and TSTr3. Notably, at 0.02 μM aTc, pronounced signal decay was observed for TSTr3, whereas TSTr1 showed little apparent signal decay. At the highest aTc concentration tested (0.2 μM), GFP output remained high over time at both cumate concentrations (Figure 3C, 3D, S7). These observations indicated that both CN and inducer concentration influenced the persistence of the reporter signal, with high aTc induction associated with sustained fluorescence and reduced apparent signal decay.

The effect of CN, aTc, and cumate concentrations on signal decay rate and fraction were quantified with a 2^3^ full factorial design. Based on the preceding observations, aTc concentrations of 0.01 (−1) and 0.02 μM (+1) and cumate concentrations of 10 (−1) and 100 μM (+1) were selected as factor levels, as these conditions allowed measurable signal decay while retaining differences in decay behavior. This resulted in eight experimental combinations (Table S7; Figure S8A). For the signal decay fraction, the medium-copy-number construct exhibited a higher decay fraction than the high-copy-number construct (Figure 3E). Regression analysis showed significant negative effects of CN, aTc, and cumate concentration on the signal decay fraction. Increasing CN, aTc concentration, and cumate concentration decreased the decay fraction, with CN showing the largest regression coefficient in magnitude (Figure S8B). aTc exerted the strongest negative effect on the decay rate constant *K*, with higher aTc concentrations slowing down apparent signal decay (Figure 3F; Figure S8C). This effect may reflect a shift in the balance between reporter production and degradation, as increased toehold switch RNA expression at higher aTc concentrations could sustain reporter production relative to degradation and result in more persistent GFP fluorescence. Together, these results indicated that CN and inducer concentrations could be used to tune the extent and rate of apparent signal decay. Based on its high ON/OFF ratio and more complete signal decay, TSTr3 was selected as the optimized module for subsequent integration with the TF-based sensing module.

### 3.4 TF–TS coupling expands the dynamic range of pyruvate sensing

To test the TF–TS hybrid architecture for metabolite sensing, we used pyruvate as a model analyte and first characterized a conventional PdhR-based pyruvate biosensor (Pyr-GFP), which showed a concentration-dependent GFP response to pyruvate (Figure S9). Based on our previous finding that reducing RBS strength can decrease basal expression and improve the dynamic range of PdhR-based biosensor(*27*), we additionally constructed Pyr-GFP (RBS33) containing a weaker RBS. We then compared two hybrid constructs, in which P_pdh_ promoter controlled either toehold switch RNA or trigger RNA, resulting in Pyr-toehold and Pyr-trigger, respectively (Figure 4A). The inducibility of the Pyr-trigger and Pyr-toehold constructs was further confirmed by time-course measurements under their respective inducer conditions (Figure S10).

Dose–response analysis revealed distinct differences in basal expression, maximum output, and dynamic range among these constructs (Figure 4B; Table S8). The conventional Pyr-GFP construct exhibited the highest maximum fluorescence but also substantial basal fluorescence, resulting in a limited dynamic range of 5.59 ± 0.09. Reducing the RBS strength in Pyr-GFP (RBS33) markedly decreased basal expression by 36.2-fold relative to Pyr-GFP, but also substantially reduced the maximum output, resulting in a dynamic range of 12.60 ± 0.24, corresponding to only a 2.2-fold increase in dynamic range over Pyr-GFP. The hybrid biosensors resulted in drastically lower basal fluorescence, with Pyr-trigger and Pyr-toehold reducing basal fluorescence approximately 503-fold and 210-fold, respectively. Pyr-trigger exhibited a dynamic range of 72.9 ± 38.2, representing a 13.04-fold increase over Pyr-GFP, while Pyr-toehold retained a substantially higher maximum, resulting in the highest dynamic range of 249.72 ± 19.64, 44.7-fold higher than Pyr-GFP and 3.4-fold higher than Pyr-trigger. Coupling PdhR-mediated pyruvate sensing to the toehold switch module effectively reduced basal reporter expression, expanding the dynamic range of the pyruvate biosensor.

### 3.5 Separated regulatory layers enable temporal control of biosensor activation

The Pyr-toehold biosensor combines transcriptional metabolite sensing with RNA-mediated translational control. Specifically, pyruvate-responsive P_pdh_ controls toehold switch RNA expression, whereas the cumate-responsive system controls trigger RNA expression. Interaction between the two RNAs activates reporter translation, thereby forming a dual-input AND-gate architecture. We therefore examined how the timing of pyruvate and cumate addition affected the onset and magnitude of the biosensor response.

Timing the addition of trigger induction revealed dynamics of pyruvate consumption in the biosensor. Addition of cumate at either 0, 2, 4, or 6 h during growth of Pyr-toehold on 10 mM pyruvate resulted in GFP activation, with the strongest response observed when cumate was added at 0 h. Delayed cumate addition gradually reduced the GFP signal expression level (Figure 5A). As pyruvate was added at the beginning of the experiment, its consumption during cell growth may have reduced P_pdh_ activation over time, thereby decreasing the amount of toehold switch RNA available for subsequent trigger-mediated activation. Next, we pre-induced trigger RNA expression with cumate and introduced pyruvate at different time points. The onset of GFP signal depended on the timing of pyruvate induction (Figure 5B), demonstrating that the timing of inducer additions modulates the onset and magnitude of Pyr-toehold activation.

**Figure 5.**
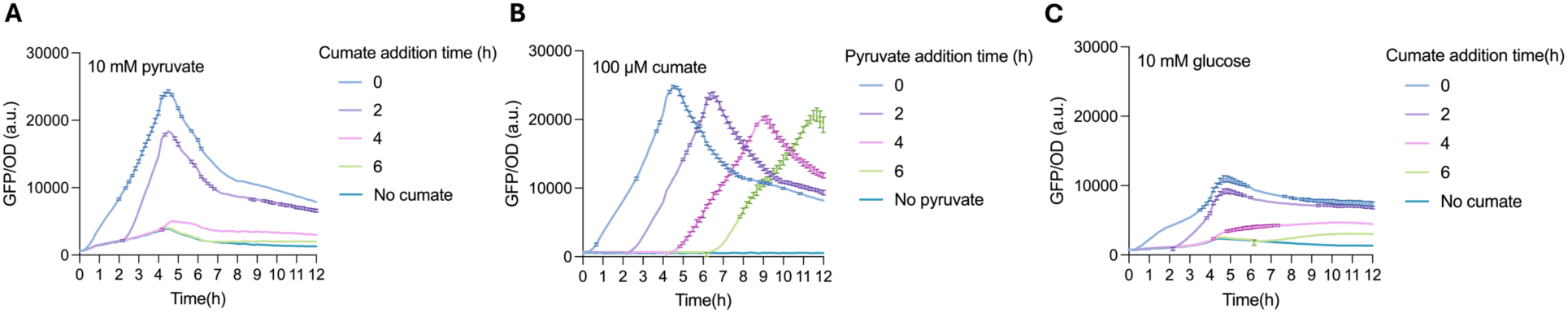
Temporal response of the Pyr-toehold biosensor to timed inducer addition. (A) GFP fluorescence of the Pyr-toehold biosensor cultured with 10 mM pyruvate, with 100 μM cumate added at 0, 2, 4, or 6 h. (B) GFP fluorescence of the Pyr-toehold biosensor cultured with 100 μM cumate, with 10 mM pyruvate added at 0, 2, 4, or 6 h. (C) GFP fluorescence of the Pyr-toehold biosensor cultured with 10 mM glucose, with 100 μM cumate added at 0, 2, 4, or 6 h.

Last, the response of the biosensor was assessed with glucose as a physiologically relevant carbon source (*33, 43*). Low GFP signal in the absence of cumate addition confirmed that PdhR-mediated production of toehold switch RNA alone was insufficient to activate the reporter. Addition of cumate at 0, 2, 4, or 6 h induced GFP expression, although the response was lower than that observed with exogenous pyruvate (Figure 5C). The lower response was associated with the higher cell density observed under glucose conditions (Figure S11). Overall, these results demonstrate that the Pyr-toehold biosensor integrates metabolite sensing and RNA-level translational control as two temporally separable regulatory layers, allowing the timing of reporter activation to be modulated through either input.

## 4. Discussion

Genetically encoded biosensors based on TFs have been widely used for the detection of small-molecule metabolites, but their regulatory performance is often limited by the direct coupling of ligand sensing and reporter expression within a single transcriptional layer, potentially resulting in high basal expression and limited dynamic range(*18*). To address this, we introduced a toehold switch system as an additional RNA-level regulatory layer. The dual-input system enabled independent control of toehold switch and trigger RNA expression and produced AND-gate behavior with tunable fluorescence output by adjusting the concentration of aTc and cumate. Compared with conventional AND-gate architectures that integrate multiple inputs primarily at the transcriptional level (*44, 45*), this configuration separates transcriptional input control from translational activation. The *E. coli* Marionette MG1655 background further facilitates this modular design because the orthogonal transcriptional regulators required to control the two RNA components are already encoded in the host(*39*), avoiding the need to introduce additional TF expression cassettes as required in some previously reported tunable genetic devices(*30*). The residual GFP signal observed with aTc alone could arise from basal expression of the cumate-controlled trigger RNA, incomplete repression of translation by the unbound TS, or a combination of these effects. Consequently, promoter leakiness and intrinsic toehold switch leakage should both be considered when selecting regulatory components, particularly when translating this architecture to other TFs or inducible systems with different basal activities.

Optimization of the TSTr-series constructs further demonstrated that tuning plasmid CN and RBS strength could improve the balance between signal amplification and basal expression, thereby enhancing the ON/OFF ratio. CN and RBS strength jointly influenced ON- and OFF-state fluorescence, such that increasing expression capacity could enhance induced output while simultaneously increasing background expression (Figure 2). This is consistent with their roles in determining genetic dosage and translation efficiency(*19, 46*). Previous studies have similarly shown that stronger RBSs can increase both maximal reporter output and basal expression(*47*). Similarly, increasing plasmid copy number increases the abundance of the genetic components and can consequently increase basal expression(*48*). This highlights that maximizing reporter output does not necessarily maximize biosensor performance; expression strength needs to be balanced against basal expression to achieve a high ON/OFF ratio. At the same time, reporter signal decay depended on plasmid CN and inducer concentrations (Figure 3), highlighting their influence on temporal responsiveness of the TSTr system. GFP fluorescence dynamics reflect the combined effects of protein synthesis, maturation, degradation, and growth dilution(*49, 50*). Higher CN or stronger induction can increase reporter synthesis relative to cellular degradation capacity, potentially slowing signal decay when GFP production approaches or exceeds its effective degradation rate(*49*). Therefore, the medium-copy-number, strong-RBS configuration, which achieved the highest ON/OFF ratio (Figure 2) and a favorable decay fraction (Figure 3), was selected for hybrid biosensor construction.

Coupling this optimized toehold switch module with PdhR sensing system demonstrated the advantage of layered regulation for metabolite sensing. Because toehold switches respond to complementary RNA triggers rather than directly recognizing small molecules, they can connect TF-mediated metabolite recognition to programmable RNA-level signal processing. Both hybrid architectures altered the response relative to direct TF-mediated reporter expression, with Pyr-toehold producing the largest increase in dynamic range (Figure 4). This improvement was primarily associated with strong suppression of basal reporter expression, demonstrating how an additional translational control layer can increase separation between OFF and ON states. However, Pyr-toehold also showed lower maximum fluorescence than the conventional PdhR biosensor. This trade-off may reflect the translational constraints of the toehold switch architecture, in which the RBS and start codon are sequestered within a designed RNA structure and must be released by trigger-mediated strand displacement before translation can initiate. The longer and more structured 5′ UTR may reduce ribosome accessibility and translation initiation efficiency compared with a conventional RBS(*51*), while incomplete or kinetically limited switch–trigger interactions may further restrict the fraction of switch transcripts that reach the fully activated state(*29*). When pyruvate is supplied as the sole carbon source, its availability may decrease during cultivation, potentially limiting sustained PdhR-mediated transcription. The secondary structure of the toehold switch may confer greater RNA stability than the trigger RNA by reducing accessibility to RNases(*29, 52*). This difference in RNA stability could partly explain the improved performance of the Pyr-toehold architecture. Overall, these results show that introducing an RNA regulatory layer can expand biosensor dynamic range, particularly through suppression of basal output, while introducing a trade-off with absolute signal amplitude.

The separation of the two regulatory layers also resulted in temporally dependent biosensor activation. Delayed cumate addition following pyruvate induction progressively reduced the maximum GFP response (Figure 5A), potentially due to reduced availability of functional toehold switch RNA over time as a result of RNA degradation and growth-associated dilution, although RNA abundance was not directly measured(*53*). Conversely, when trigger RNA was pre-induced, subsequent pyruvate addition shifted the onset of GFP activation according to the timing of pyruvate induction (Figure 5B). This indicates that, when cumate induction maintains a sustained pool of trigger RNA in the cells, the system can rapidly respond to the subsequent availability of pyruvate as a carbon source. These results demonstrate that the Pyr-toehold architecture provides not only dual-input logic but also a means of exploiting the temporal availability of each regulatory input. More broadly, this TF–TS hybrid architecture provides a modular framework in which metabolite recognition and downstream signal processing can be engineered as partially separable functions. It offers opportunities to tune biosensor dynamic range, basal expression, and temporal response.

## 5. Conclusion

In this study, we developed a modular TF–TS hybrid architecture that introduces an additional RNA-level regulatory layer into a whole-cell biosensor for pyruvate sensing. We first established a dual-input TSTr system with independently regulated switch and trigger RNA expression. Through DoE-guided optimization, TSTr3 achieved the highest ON/OFF ratio of 33.70 ± 2.36, representing a 19.8-fold increase relative to the weakest-performing construct. Signal decay analysis further showed plasmid CN and inducer concentrations affected the temporal responsiveness of the system. The optimized module was subsequently integrated with a PdhR-based pyruvate biosensor. The resulting Pyr-toehold biosensor achieved a dynamic range of 249.72 ± 19.64, representing a 44.7-fold increase in dynamic range and a 210-fold reduction in basal expression compared with the conventional PdhR-based architecture. Finally, temporal induction experiments demonstrated that the two regulatory layers could influence the timing and magnitude of biosensor activation. This architecture could be further extended to other ligand-responsive sensing systems, offering a promising strategy for engineering biosensors with improved dynamic range and tunable response dynamics.

## 6. CRediT authorship contribution statement

Zihan Gao: Writing – original draft, methodology, investigation, formal analysis, data curation. Maria Suarez-Diez: Writing – review & editing, supervision, validation. Pieter Candry: Writing – review & editing, supervision, validation.

## 7. Acknowledgements

We acknowledge Jelmar de Vries for valuable discussions and suggestions regarding the use of the *E. coli* Marionette strain. We thank Jenny Bakker for providing plasmid pSEVAb86 and Silvia Rodríguez Marcos for experimental material support. We acknowledge the assistance of Güniz Özer-Bergman during the initial discussions. Zihan Gao was supported by the China Scholarship Council (CSC) under Grant No. 202304910065. Pieter Candry was supported by the Dutch Research Council (NWO) Veni Talent Programme (File No. 21027) and the Dutch Sectorplan Bèta-II – Biology. Schematics were created with BioRender.

## 8. Data availability statement

The data supporting the findings of this study are available on Zenodo at https://doi.org/10.5281/zenodo.24470413.

## 9. Appendix A. Supplementary data

The Supporting Information is available free of charge.

