## Supplementary Material for "Integration with a toehold switch expands the dynamic range of a transcription factor-based whole-cell pyruvate biosensor"

**Table S1. Strains used in this study.**
TS, toehold switch only; Tr, trigger RNA; TSTr, co-expression of the toehold switch and trigger RNA; Pyr-trigger, $P_{pdh}$-driven trigger RNA expression; Pyr-toehold, $P_{pdh}$-driven toehold switch RNA expression.

| Strain | Description | Reference |
|---|---|---|
| *E. coli* DH5α | Cloning strain | Lab stock |
| *E. coli* BL21 (DE3) | Protein expression strain | Lab stock |
| *E. coli* Marionette MG1655 | MG1655-derived strain engineered with 12 orthogonal transcription factor sensors genomically integrated into the chromosome | *(1)* |
| TS | *E. coli* BL21 (DE3) containing pUC-TS | This study |
| TS-Tr | *E. coli* BL21 (DE3) containing pUC-TS-Tr | This study |
| TS (RBS33) | *E. coli* BL21 (DE3) containing pUC-TS (RBS33) | This study |
| TS (RBS33)-Tr | *E. coli* BL21 (DE3) containing pUC-TS (RBS33)-Tr | This study |
| $P_{tet}$_toehold_gfp_ $P_{cymRC}$_trigger | *E. coli* Marionette MG1655 containing pSEVA-$P_{tet}$-TS-gfp-$P_{cymRC}$-Tr | This study |
| TSTr1 | *E. coli* Marionette MG1655 containing pUC-TSTr-1 | This study |
| TSTr2 | *E. coli* Marionette MG1655 containing pUC-TSTr (RBS33)-2 | This study |
| TSTr3 | *E. coli* Marionette MG1655 containing pSEVA-TSTr-3 | This study |
| TSTr4 | *E. coli* Marionette MG1655 containing pSEVA-TSTr (RBS33)-4 | This study |
| Pyr-GFP | *E. coli* Marionette MG1655 containing Pyr-GFP | This study |
| Pyr-GFP (RBS33) | *E. coli* Marionette MG1655 containing Pyr-GFP (RBS33) | This study |
| Pyr-trigger | *E. coli* Marionette MG1655 containing Pyr-trigger | This study |
| Pyr-toehold | *E. coli* Marionette MG1655 containing Pyr-toehold | This study |

**Table S2. Plasmids used in this study.**

P represents promoter, RBS represents ribosome binding site, and t represents the B0010 terminator.

| Plasmid | Description | Reference |
|---|---|---|
| pUC19 | Vector, ColE1 origin, high copy number, $Amp^R$ | Lab stored |
| pSEVAb86 | pSEVAb (SEVA 3.1) vector, p15A origin, medium copy number, $Apr^R$ | Lab stored |
| pUC-TS | pUC19 vector carrying $P_{J115}$-RBS-toehold switch controlling gfp-ASV-t | This study |
| pUC-TS-Tr | pUC19 vector carrying $P_{J115}$-RBS-toehold switch controlling gfp-ASV-t, together with $P_{J100}$-trigger-t | This study |
| pUC-TS (RBS33) | pUC19 vector carrying $P_{J115}$-RBS33-toehold switch controlling gfp-ASV-t | This study |
| pUC-TS (RBS33)-Tr | pUC19 vector carrying $P_{J115}$-RBS33-toehold switch controlling gfp-ASV-t, together with $P_{J100}$-trigger-t | This study |
| pSEVA-$P_{tet}$-TS-gfp-$P_{cymRC}$-Tr | pSEVAb86 vector carrying $P_{tet}$-RBS-toehold switch regulating gfp-t, together with a $P_{cymRC}$-driven trigger RNA-t | This study |
| pUC-TSTr-1 | pUC19 vector carrying $P_{tet}$-RBS-toehold switch regulating gfp-ASV-t, together with a $P_{cymRC}$-trigger-t | This study |
| pUC-TSTr (RBS33)-2 | pUC19 vector carrying $P_{tet}$-RBS33-toehold switch regulating gfp-ASV-t, together with a $P_{cymRC}$-trigger-t | This study |
| pSEVA-TSTr-3 | pSEVAb86 vector carrying $P_{tet}$-RBS-toehold switch regulating gfp-ASV-t, together with a $P_{cymRC}$-trigger-t | This study |
| pSEVA-TSTr (RBS33)-4 | pSEVAb86 vector carrying $P_{tet}$-RBS33-toehold switch regulating gfp-ASV-t, together with a $P_{cymRC}$-trigger-t | This study |
| Pyr-GFP | pSEVAb86 vector carrying $P_{pdh}$-RBS regulating gfp-ASV-t | This study |
| Pyr-GFP (RBS33) | pSEVAb86 vector carrying $P_{pdh}$-RBS33 regulating gfp-ASV-t | This study |
| Pyr-trigger | pSEVAb86 vector carrying $P_{tet}$-RBS-toehold switch regulating gfp-ASV-t, together with a $P_{pdh}$-trigger-t | This study |
| Pyr-toehold | pSEVAb86 vector carrying $P_{pdh}$-RBS-toehold switch regulating gfp-ASV-t, together with a $P_{cymRC}$-trigger-t | This study |

**Table S3. Genetic parts and sequences used in this study.**
The -35 and -10 boxes are underlined. Operators are shown in red.

| Part name | Description | DNA sequence (5′–3′) |
|---|---|---|
| $P_{J115}$ (J23115) | Constitutive promoter | TTTATAGCTAGCTCAGCCCTTGGTACAATGCTAGC |
| $P_{J100}$ (J23100) | Constitutive promoter | TTGACGGCTAGCTCAGTCCTAGGTACAGTGCTAGC |
| $P_{pdh}$ | Inducible promoter | TGGACATAAGGTGAATACTTTGTTACTTTAGCGTCACAGACATGAAATTGGTAAGACCAATTGACTTCGGCAAGTGGCTTAAGACAGGAACTC |
| $P_{tet}$ | Inducible promoter | TCCCTATCAGTGATAGAGATTGACATCCCTATCAGTGATAGAGATACTGAGCACATCAGCAGGACGCACTGACC |
| $P_{cymRC}$ | Inducible promoter | AACAAACAGACAATCTGGTCTGTTTGTATTATGGAAAATTTTTCTGTATAATAGATTCAACAAACAGACAATCTGGTCTGTTTGTAT |
| RBS | Ribosome binding site | AGAGGAGA |
| RBS33 (B0033) | Ribosome binding site | TCACACAGGAC |
| TS | Toehold switch RNA | GGGCGTTAATCTCTGGCTTGCTTTATGTCTGTAAACAGAGGAGATACAGAATGAAAGCAAGCAACCTGGCGGCAGCGCAAAAGATGCGTAAA |
| TS (RBS33) | Toehold switch (RBS33) RNA | GGGCGTTAATCTCTGGCTTGCTTTATGTCTGTAAACTCACACAGGACTACAGAATGAAAGCAAGCAACCTGGCGGCAGCGCAAAAGATGCGTAAA |
| Tr | Trigger RNA | GGGTCATGACTGGGACACGCCAGTCATGAGAATACAGACATAAAGCAAGCCAGAGATTAACGAAG |
| *sfgfp* | Reporter gene | ATGCGTAAAGGCGAAGAGCTGTTCACGGGCGTAGTTCCGATTCTGGTCGAGCTGGACGGCGATGTGAACGGTCATAAGTTTAGCGTTCGCGGTGAAGGTGAGGGCGACGCGACCAACGGCAAACTGACCCTGAAGTTCATCTGCACCACCGGTAAACTGCCGGTGCCTTGGCCGACCTTGGTGACGACGTTGACGTATGGCGTGCAGTGTTTTGCGCGTTATCCGGACCACATGAAACAACACGATTTCTTCAAATCTGCGATGCCGGAGGGTTACGTCCAGGAGCGTACCATTTCCTTCAAGGATGATGGCTACTACAAAACTCGCGCAGAGGTTAAGTTTGAAGGTGACACGCTGGTCAATCGTATCGAATTGAAGGGTATCGACTTTAAAGAGGATGGTAACATTCTGGGCCATAAACTGGAGTATAACTTCAACAGCCATAATGTTT |

| | | |
|---|---|---|
| | | ACATTACGGCAGACAAGCAAAAGAACGGC<br>ATCAAGGCCAATTTCAAGATTCGCCACAAT<br>GTTGAGGACGGTAGCGTCCAACTGGCCGAC<br>CATTACCAGCAGAACACCCCAATTGGTGAC<br>GGTCCGGTTTTGCTGCCGGATAATCACTATC<br>TGAGCACCCAAAGCGTGCTGAGCAAAGATC<br>CGAACGAAAAACGTGATCACATGGTCCTGC<br>TGGAATTTGTGACCGCTGCGGGCATCACCC<br>ACGGTATGGATGAACTGTACAAA |
| ASV | Degradation tag | AGGCCAGCAGCAAACGACGAAAACTACGCT<br>GCATCAGTT |
| B0010 | Terminator | CCAGGCATCAAATAAAACGAAAGGCTCAGT<br>CGAAAGACTGGGCCTTTCGTTTTATCTGTTG<br>TTTGTCGGTGAACGCTCTC |
| *pdhR* | Transcriptional repressor from *E. coli* MG1655 | ATGGCCTACAGCAAAATCCGCCAACCAAAA<br>CTCTCCGATGTGATTGAGCAGCAACTGGAG<br>TTTTTGATCCTCGAAGGCACTCTCCGCCCGG<br>GCGAAAAACTCCCACCGGAACGCGAACTGG<br>CAAAACAGTTTGACGTCTCCCGTCCCTCCTT<br>GCGTGAGGCGATTCAACGTCTCGAAGCGAA<br>GGGCTTGTTGCTTCGTCGCCAGGGTGGCGG<br>CACTTTTGTCCAGAGCAGCCTATGGCAAAG<br>CTTCAGCGATCCGCTGGTGGAGCTGCTCTCC<br>GACCATCCTGAGTCACAGTATGACTTGCTC<br>GAAACACGACACGCCCTGGAAGGTATCGCC<br>GCTTATTACGCCGCGCTGCGTAGTACCGAT<br>GAAGACAAGGAACGCATCCGTGAACTCCAC<br>CACGCCATAGAGCTGGCGCAGCAGTCTGGC<br>GATCTGGACGCGGAATCAAACGCCGTACTC<br>CAGTATCAGATTGCCGTCACCGAAGCGGCC<br>CACAATGTGGTTCTGCTTCATCTGCTAAGGT<br>GTATGGAGCCGATGTTGGCCCAGAATGTCC<br>GCCAGAACTTCGAATTGCTCTATTCGCGTCG<br>CGAGATGCTGCCGCTGGTGAGTAGTCACCG<br>CACCCGCATATTTGAAGCGATTATGGCCGG<br>TAAGCCGGAAGAAGCGCGCGAAGCATCGC<br>ATCGCCATCTGGCCTTTATCGAAGAAATTTT<br>GCTCGACAGAAGTCGTGAAGAGAGCCGCCG<br>TGAGCGTTCTCTGCGTCGTCTGGAGCAACG<br>AAAGAATTAG |

**Table S4. Oligonucleotides used in this study.**

| Primer name | Sequence (5'-3') | Usage |
|---|---|---|
| 1_verify_fwd | GTTGTGTGGAATTGTGAGCGG | Verification of pUC-series plasmids |
| 2_verify_rev | TCCCAGTCACGACGTTGTAAAAC | |
| 3_verify_fwd | GCGCGCCCAGCTGTCTAG | Verification of pSEVA-series plasmids |
| 4_verify_rev | CGTAATTATTGGGGACCCCTGG | |
| 5_gfp_ASV_fwd | CATGCCTGCAGGTCGACTCTGAGAGCGTTCACCGACAAACAACAGATAAAACGAAAGGCCCAGTCTTTCGACTGAGCCTTTCGTTTTATTTGATGCCTGGTCAAACTGATGCAGCGTAGTTTTCGTCGTTTGCTGCTGGCCTTTTGTACAGTTCATCCATACCGTG | Construction of pUC-TS |
| 6_gfp_ASV_rev | CGAGCTCGGTACCCGGGGATTTTATAGCTAGCTCAGCCCTTGGTACAATGCTAGCGGGCGTTAATCTCTGGCTTGCTTTATGTCTGTAAACAGAGGAGATACAGAATGAAAGCAAGCAACCTGGCGGCAGCGCAAAAGATGCGTAAAATGCGTAAAGGCGAAGAGCT | |
| 7_backbone_fwd | ATCCCCGGGTACCGAGCTCGAATTC | |
| 8_backbone_rev | AGAGTCGACCTGCAGGCATG | |
| 9_backbone_fwd | CCGGGTACCGAGCTCGAATTC | Amplification of the pUC backbone for construction of pUC-TSTr |
| 10_backbone_rev | TTTATAGCTAGCTCAGCCCTTG | |
| 11_RBS33_fwd | GTCCTGTGTGAGTTTACAGACATAAAGCAAGCCAGAGATTAACGC | Replacement of the original RBS with RBS33 in pUC-TS |
| 12_RBS33_rev | TCTGTAAACTCACACAGGACTACAGAATGAAAGCAAGCAACCTGG | |
| 13_gfp_fwd | CATGATTACGCCAAGCTTGCATG | Construction of pSEVA-Ptet-TS-gfp-PcymRC-Tr plasmid |
| 14_gfp_rev | ATGCGTAAAGGCGAAGAGCTGTTC | |
| 15_backbone_fwd | GAACAGCTCTTCGCCTTTACGC | |
| 16_backbone_rev | CATGCAAGCTTGGCGTAATCATG | |
| 17_backbone_fwd | ATCCCCGGGTACCGAGCTC | Amplification of the plasmid backbone for construction of pUC-TSTr-1 |
| 18_backbone_rev | ATGCGTAAAGGCGAAGAGCTG | |
| 19_TSTr_fwd | GAATTCGCGGCCGCTTCTAGAGGTTGTGTGGAATTGTGAGCGG | Construction of pSEVA-TSTr-3 |
| 20_TSTr_rev | CTGCAGCGGCCGCTACTAGTATCCCAGTCACGACGTTGTAAAAC | |
| 21_backbone_fwd | TACTAGTAGCGGCCGCTGCAG | |
| 22_backbone_rev | CTCTAGAAGCGGCCGCGAATTC | |
| 23_Ppdh_fwd | GAACAGCTCTTCGCCTTTAC | |

| | | |
|---|---|---|
| 24_Ppdh_rev | CTGCAGCGGCCGCTACTAGTA | Construction of Pyr-GFP |
| 25_backbone_fwd | TACTAGTAGCGGCCGCTGCAG | |
| 26_gfp_rev | ATGCGTAAAGGCGAAGAGCTGTTC | |
| 27_RBS33_fwd | GTCCTGTGTGAGTTTGAAGTCAATTGGTCTTACC | Replacement of the original RBS with RBS33 in Pyr-GFP |
| 28_RBS33_rev | ACTTCAAACTCACACAGGACTACAGAAATGCGTAAAGGC | |
| 29_fwd | GTGACGCTAAAGTAACAAAGTATTCACCTTATGTCCATACAGGTTTTACAGGTTTTGATTGAAATCATGAAACTG | Construction of Pyr-toehold |
| 30_rev | CTTTGTTACTTTAGCGTCACAGACATGAAATTGGTAAGACCAATTGACTTCGGGCGTTAATCTCTGGCTTG | |
| 31_fwd | TGAATACTTTGTTACTTTAGCGTCACAGACATGAAATTGGTAAGACCAATTGACTTCGGGTCATGACTGGGACACGC | Construction of Pyr-trigger |
| 32_rev | CTAAAGTAACAAAGTATTCACCTTATGTCCATACAGGTTTGTCTACATTTGTGCATAGTTACAACTTTGAAAC | |

**Table S5. Performance characteristics of TSTr-series constructs from the $2^2$ full factorial design.**

| Construct | CN | RBS | ON (a.u.) | OFF (a.u.) | ON/OFF |
|---|---|---|---|---|---|
| TSTr1 | 1 | 1 | 41411.42 ± 1267.96 | 3959.31 ± 101.59 | 10.46 ± 0.27 |
| TSTr2 | 1 | -1 | 2822.18 ± 154.82 | 731.74 ± 58.78 | 3.88 ± 0.45 |
| TSTr3 | -1 | 1 | 5280.35 ± 194.80 | 157.07 ± 8.98 | 33.70 ± 2.36 |
| TSTr4 | -1 | -1 | 127.66 ± 22.96 | 77.16 ± 20.92 | 1.71 ± 0.35 |

CN, copy number; RBS, ribosome binding site. The high-copy-number ColE1 plasmid and the strong RBS were coded as +1, whereas the medium-copy-number p15A plasmid and the weak RBS33 were coded as −1. ON/OFF was calculated for each biological replicate and is reported as mean ± SD.

**Table S6. Regression analysis of TSTr-series construct performance.**

Dependent variable: ON/OFF

| Term | Estimate | *P* value | Significance |
|---|---|---|---|
| Intercept | 12.44 | $4.57 \times 10^{-10}$ | *** |
| CN | -5.27 | $3.99 \times 10^{-7}$ | *** |
| RBS | 9.64 | $3.45 \times 10^{-9}$ | *** |
| CN × RBS | -6.35 | $9.25 \times 10^{-8}$ | *** |

$Y_{(ON/OFF)} = 12.44 - 5.27\ CN + 9.64\ RBS - 6.35\ (CN \times RBS)$

$R^2 = 0.9939$; Adjusted $R^2 = 0.9916$; $P$ value $= 3.484 \times 10^{-9}$

Dependent variable: ON (a.u.)

| Term | Estimate | *P* value | Significance |
|---|---|---|---|
| Intercept | 12410.4 | $2.90 \times 10^{-12}$ | *** |
| CN | 9706.4 | $2.06 \times 10^{-11}$ | *** |
| RBS | 10935.5 | $7.96 \times 10^{-12}$ | *** |
| CN × RBS | 8359.1 | $6.79 \times 10^{-11}$ | *** |
| $Y_{(ON)} = 12410.4 + 9706.4\ CN + 10935.5\ RBS + 8359.1\ (CN \times RBS)$ | | | |
| $R^2 = 0.9990$; Adjusted $R^2 = 0.9987$; *P* value = $2.272 \times 10^{-12}$ | | | |

Dependent variable: OFF (a.u.)

| Term | Estimate | *P* value | Significance |
|---|---|---|---|
| Intercept | 1231.3 | $1.66 \times 10^{-12}$ | *** |
| CN | 1114.2 | $3.68 \times 10^{-12}$ | *** |
| RBS | 826.9 | $3.98 \times 10^{-11}$ | *** |
| CN × RBS | 786.9 | $5.91 \times 10^{-11}$ | *** |
| $Y_{(OFF)} = 1231.3 + 1114.2\ CN + 826.9\ RBS + 786.9\ (CN \times RBS)$ | | | |
| $R^2 = 0.9991$; Adjusted $R^2 = 0.9987$; *P* value = $1.883 \times 10^{-12}$ | | | |

Dependent variable: Decay fraction

| Term | Estimate | *P* value | Significance |
|---|---|---|---|
| Intercept | 0.5638 | $< 2 \times 10^{-16}$ | *** |
| CN | -0.2112 | $< 2 \times 10^{-16}$ | *** |
| aTc | -0.1225 | $4.47 \times 10^{-15}$ | *** |
| Cuma | -0.0794 | $3.78 \times 10^{-12}$ | *** |
| CN × Cuma | -0.0335 | $8.72 \times 10^{-7}$ | *** |
| CN × aTc | -0.0153 | 0.00282 | ** |
| aTc × Cuma | -0.0390 | $1.18 \times 10^{-7}$ | *** |
| $Y_{(Decay\ fraction)} = 0.5638 - 0.2112\ CN - 0.1225\ aTc - 0.0794\ Cuma - 0.0335\ (CN \times Cuma) - 0.0153\ (CN \times aTc) - 0.0390\ (aTc \times Cuma)$ | | | |
| $R^2 = 0.9956$; Adjusted $R^2 = 0.9937$; *P* value $< 2 \times 10^{-16}$ | | | |

Dependent variable: Decay rate constant *K* ($h^{-1}$)

| Term | Estimate | *P* value | Significance |
|---|---|---|---|
| Intercept | 0.4194 | $< 2 \times 10^{-16}$ | *** |
| CN | -0.0409 | $3.94 \times 10^{-6}$ | *** |
| aTc | -0.1534 | $1.50 \times 10^{-13}$ | *** |

| | | | |
|---|---|---|---|
| Cuma | -0.1085 | $1.68 \times 10^{-11}$ | *** |
| CN × Cuma | -0.0627 | $2.32 \times 10^{-8}$ | *** |
| CN × aTc | -0.0204 | 0.00265 | ** |
| aTc × Cuma | 0.0068 | 0.24545 | ns |

$Y_{(\text{Decay rate constant})}$ = 0.4194 - 0.0409 CN - 0.1534 aTc - 0.1085 Cuma - 0.0627 (CN × Cuma) - 0.0204 (CN × aTc) + 0.0068 (aTc × Cuma)

$R^2 = 0.9895$; Adjusted $R^2 = 0.9851$; $P$ value = $4.814 \times 10^{-13}$

**Table S7. Degradation kinetics of the $2^3$ full factorial design experiments.**

| Run | CN | aTc | Cumate | Decay fraction | Decay rate constant $K$ ($h^{-1}$) | Peak GFP/OD (a.u.) |
|---|---|---|---|---|---|---|
| 1 | 1 | -1 | -1 | 0.56 ± 0.02 | 0.73 ± 0.03 | 22232.41 ± 344.37 |
| 2 | 1 | -1 | 1 | 0.45 ± 0.02 | 0.46 ± 0.003 | 30055.68 ± 1048.31 |
| 3 | 1 | 1 | -1 | 0.33 ± 0.05 | 0.28 ± 0.02 | 25883.87 ± 367.09 |
| 4 | 1 | 1 | 1 | 0.06 ± 0.02 | n.a. | 38267.43 ± 806.57 |
| 5 | -1 | -1 | -1 | 0.89 ± 0.01 | 0.65 ± 0.03 | 2435.75 ± 101.91 |
| 6 | -1 | -1 | 1 | 0.84 ± 0.001 | 0.46 ± 0.01 | 3353.79 ± 48.10 |
| 7 | -1 | 1 | -1 | 0.79 ± 0.01 | 0.45 ± 0.02 | 2912.83 ± 31.18 |
| 8 | -1 | 1 | 1 | 0.58 ± 0.01 | 0.29 ± 0.01 | 4154.84 ± 14.24 |

CN, copy number. The high- and low-copy constructs were coded as +1 and −1, respectively; aTc concentrations of 0.02 and 0.01 μM were coded as +1 and −1, respectively; and cumate concentrations of 100 and 10 μM were coded as +1 and −1, respectively.

**Table S8. Best-fit parameters of the dose–response curves for the pyruvate biosensors.**

| Construct | Minimum output (a.u.) | Maximum output (a.u.) | Dynamic range (max/min) | $R^2$ |
|---|---|---|---|---|
| Pyr-GFP | 17135 ± 92.73 | 95818 ± 1114.2 | 5.59 ± 0.09 | 0.9979 ± 0.0005 |
| Pyr-GFP (RBS33) | 473.27 ± 5.69 | 5964 ± 162.25 | 12.60 ± 0.24 | 0.9814 ± 0.0016 |
| Pyr-trigger | 34.09 ± 21.79 | 1934 ± 146.67 | 72.90 ± 38.2 | 0.9993 ± 0.0005 |
| Pyr-toehold | 81.79 ± 4.61 | 20371 ± 877.52 | 249.72 ± 19.64 | 0.9975 ± 0.0002 |

Maximum and minimum output values were obtained from the fitted Hill equation ($y_{max}$ and $y_{min}$, respectively). The dynamic range ($y_{max}/y_{min}$) was calculated for each biological replicate as $y_{max}/y_{min}$ and is reported as mean ± SD. $R^2$ indicates the coefficient of determination for the Hill-curve fit.

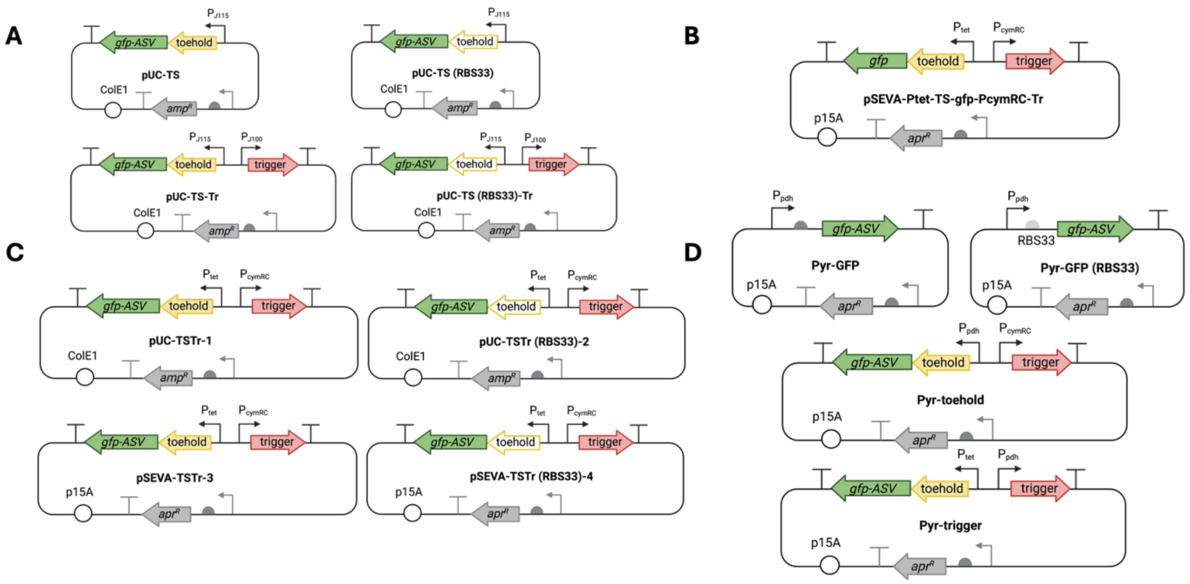


**Figure S1. Schematic overview of plasmid constructs used in this study.** (A) Toehold switch constructs used for initial functional characterization. (B) Dual-input TSTr construct for tunable induction. (C) TSTr-series constructs used for Design of Experiments (DoE)-guided optimization. (D) TF-based and hybrid TF–TS pyruvate biosensor constructs.

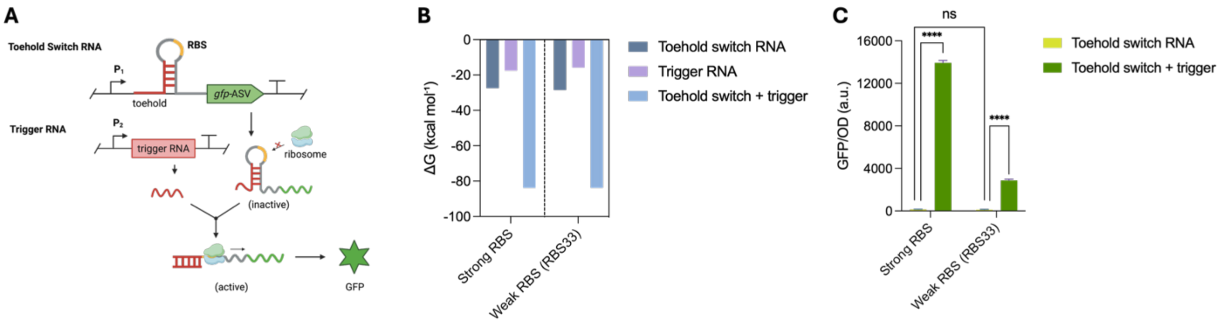


**Figure S2. Construction and validation of the toehold switch system.** (A) Schematic illustration of the toehold switch system mechanism. In the absence of trigger RNA, the RBS is sequestered within the hairpin structure, preventing translation initiation. Upon trigger RNA binding, the hairpin structure is unfolded, exposing the RBS and enabling sfGFP translation. (B) Predicted free energy (ΔG) of the toehold switch RNA, trigger RNA, and toehold switch–trigger complex. (C) GFP fluorescence characterization of toehold switch constructs containing either a strong RBS or a weak RBS (RBS33). The experiments were performed in triplicate ($n$ = 3 biologically independent samples), and results are presented as mean ± SD. Statistical significance was analyzed using Student's t-test (****$P < 0.0001$; ns, not significant).

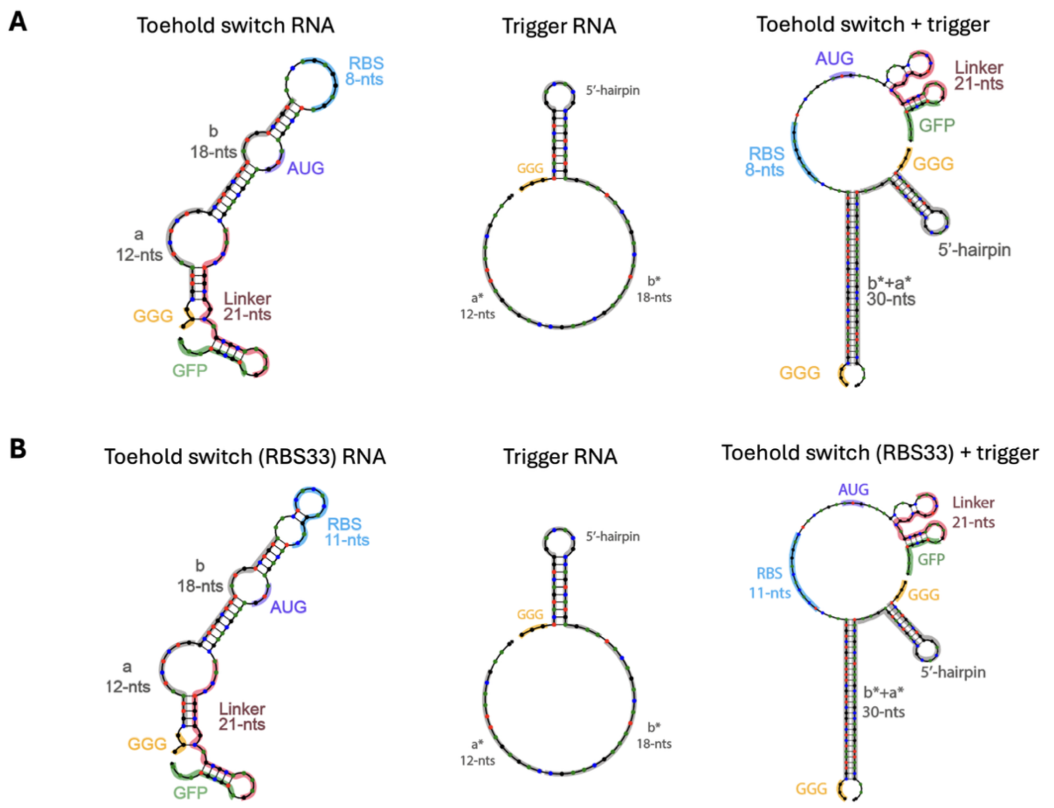


**Figure S3. Predicted secondary structures of the toehold switch variants and trigger RNA.** (A) Predicted secondary structures of the original toehold switch RNA, trigger RNA, and their interaction complex. The toehold switch RNA represses translation by sequestering the RBS and start codon within a secondary structure, thereby preventing ribosome binding and translation initiation. The a and b domains are 12 and 18 nt long, respectively, with complementary a* and b* domains present in the trigger RNA. (B) Predicted secondary structures of the RBS33 toehold switch RNA, trigger RNA, and their interaction complex.

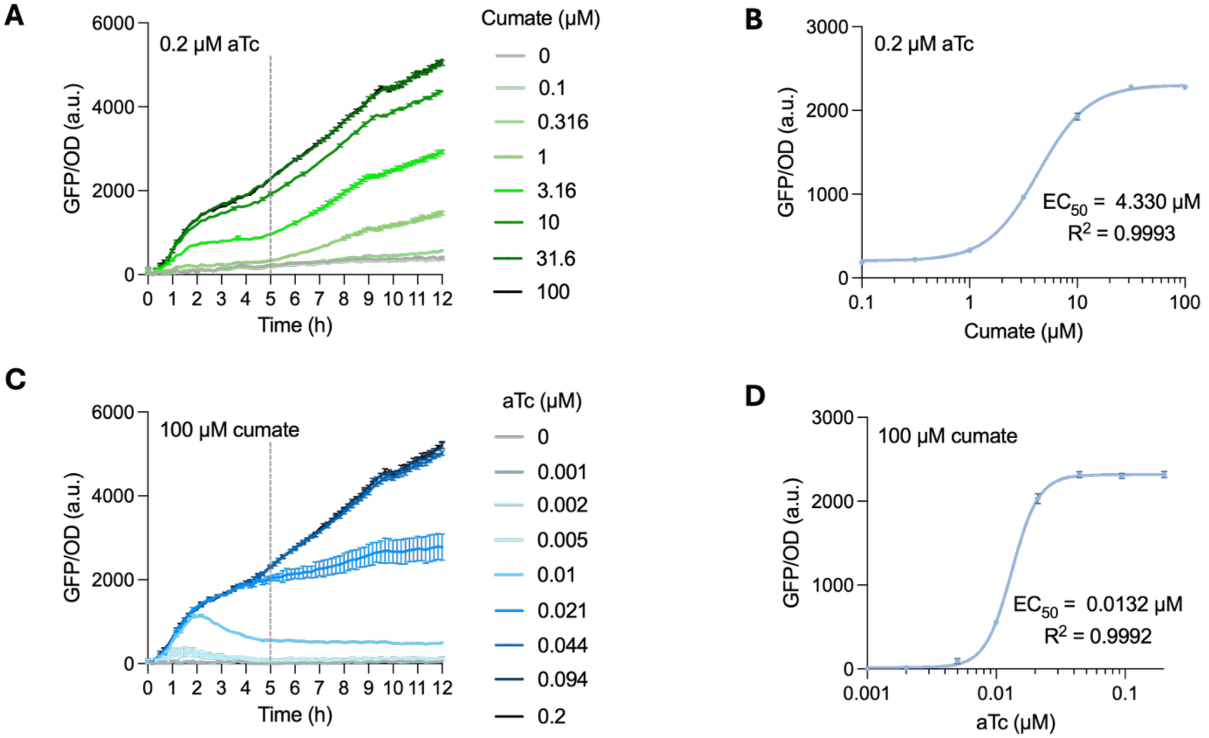


**Figure S4. Dose–response characterization of the dual-input TSTr system.** (A) Time-course GFP fluorescence under different cumate concentrations with aTc maintained at its maximum tested concentration (0.2 μM). (B) Dose–response curve of GFP fluorescence as a function of cumate concentration at 5 h. (C) Time-course GFP fluorescence under different aTc concentrations with cumate maintained at its maximum tested concentration (100 μM). (D) Dose–response curve of GFP fluorescence as a function of aTc concentration at 5 h.

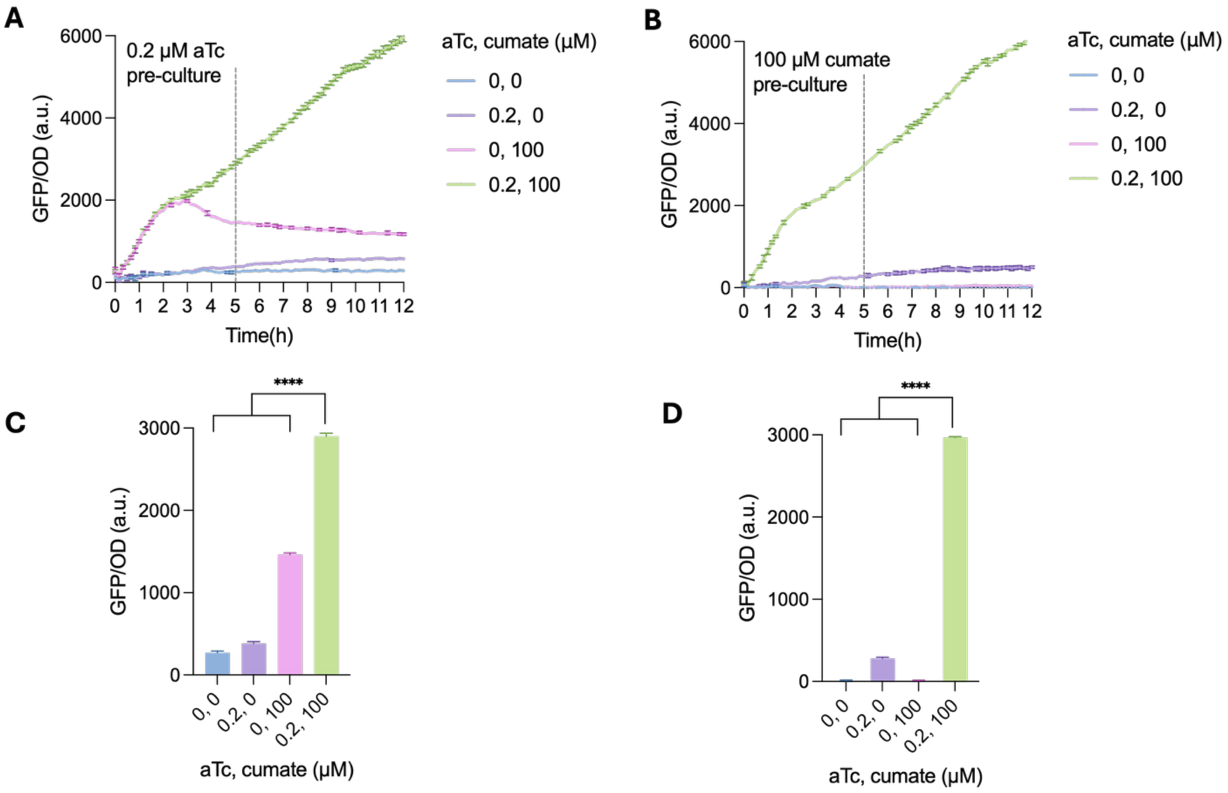


**Figure S5. Effect of pre-culture conditions on the response of the dual-input TSTr system.** (A) Time-course GFP fluorescence following pre-culture with 0.2 μM aTc and induction with different inducer conditions. (B) Time-course GFP fluorescence following pre-culture with 100 μM cumate and induction with different inducer conditions. (C) GFP fluorescence at 5 h following pre-culture with 0.2 μM aTc and induction with different inducer conditions. (D) GFP fluorescence at 5 h following pre-culture with 100 μM cumate and induction with different inducer conditions.

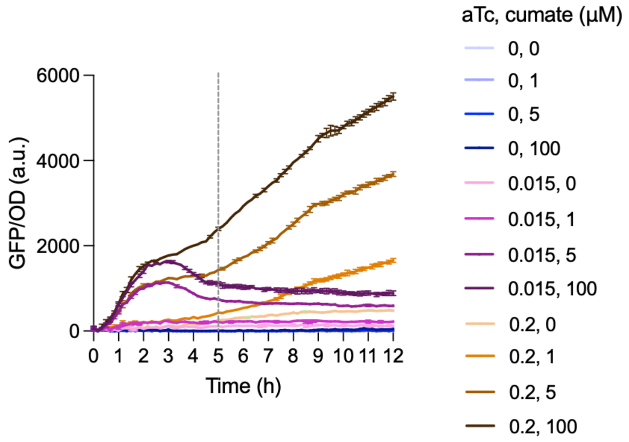


**Figure S6. Time-course characterization of the dual-input TSTr system under different aTc and cumate concentrations.**

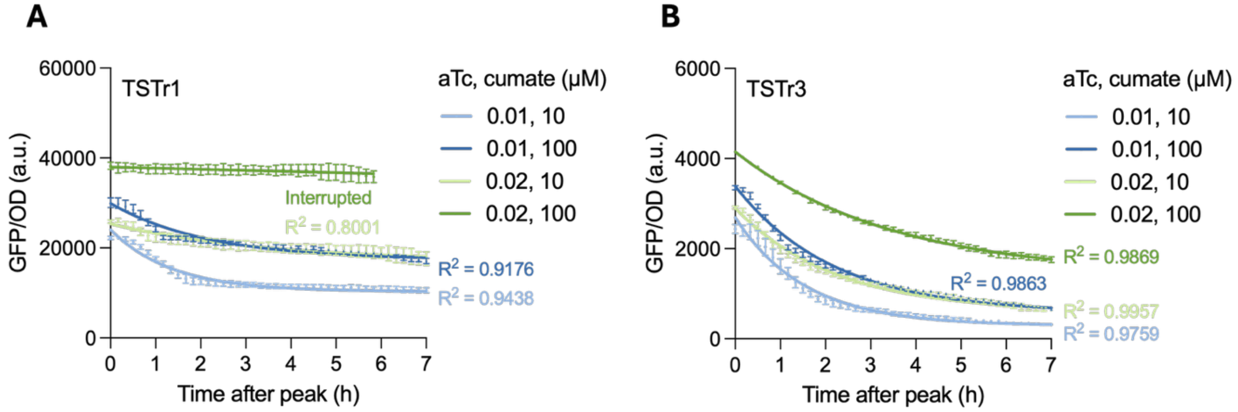


**Figure S7. One-phase decay fitting of GFP fluorescence for TSTr1 and TSTr3.** (A) Signal decay of the high-copy-number TSTr1 construct. (B) Signal decay of the medium-copy-number TSTr3 construct. GFP fluorescence decay was measured at 0.01 or 0.02 µM aTc in combination with 10 or 100 µM cumate and fitted using a one-phase decay model. Solid lines represent the fitted decay curves, and $R^2$ values indicate the goodness of fit. The 0.02 µM aTc and 100 µM cumate condition for TSTr1 showed no appreciable signal decay and was therefore not fitted.

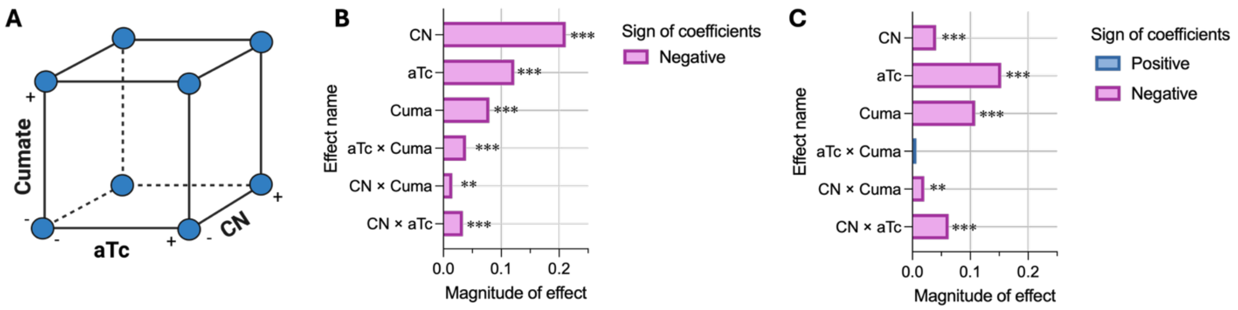


**Figure S8. Full factorial design and regression analysis of factors affecting signal decay kinetics.** (A) Schematic representation of the $2^3$ full factorial design with CN, aTc, and cumate concentration as the three factors. (B) Regression coefficients of the tested factors for the signal decay fraction. (C) Regression coefficients of the tested factors for the decay rate constant ($K$). Bars represent the estimated effects of the corresponding factors.

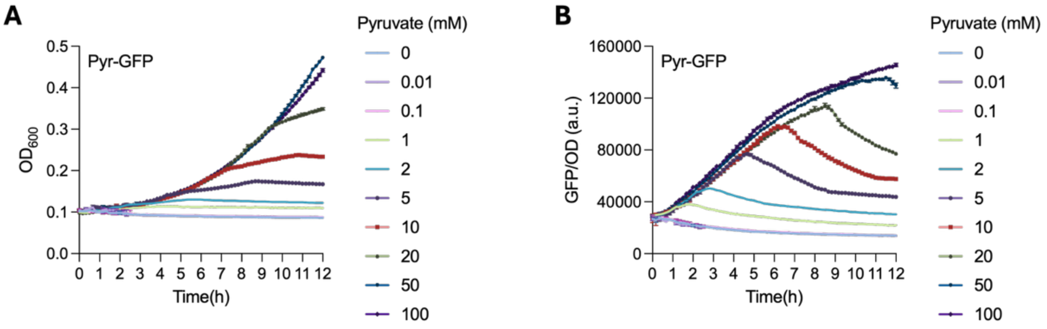


**Figure S9. Characterization of the PdhR-based pyruvate biosensor.** (A) Cell growth ($OD_{600}$) and (B) GFP fluorescence at different pyruvate concentrations over time.

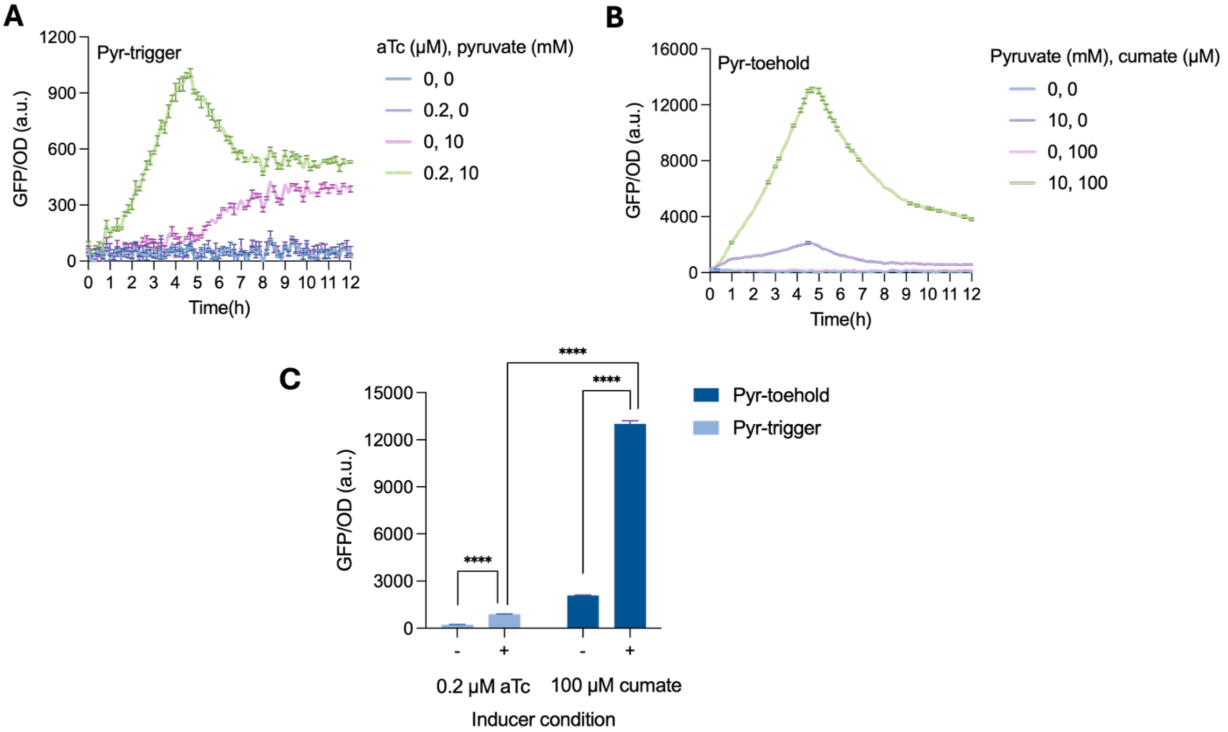


**Figure S10. Characterization of the TF–TS hybrid pyruvate biosensor.** (A) GFP fluorescence of Pyr-trigger under different aTc and pyruvate conditions. (B) GFP fluorescence of Pyr-toehold under different pyruvate and cumate conditions. (C) GFP fluorescence under the indicated inducer conditions at 5 h.

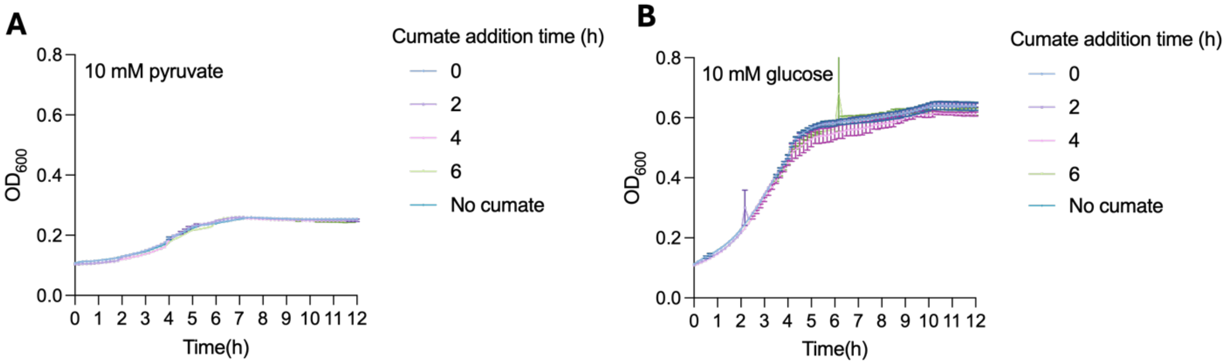


**Figure S11. Cell growth of the Pyr-toehold biosensor under different cumate addition times.** (A) Cell growth in 10 mM pyruvate. (B) Cell growth in 10 mM glucose. Transient increases in $OD_{600}$ observed at certain time points may have resulted from cell debris following cumate injection or other measurement artifacts and did not affect the overall growth profiles.

### Supplementary Methods

**Method S1. Media and stock solution preparation.**

Lysogeny broth (LB) medium contained 10 g/L tryptone, 10 g/L NaCl, and 5 g/L yeast extract. M9 minimal medium contained 3.88 g/L $K_2HPO_4$, 1.63 g/L $NaH_2PO_4$, 2 g/L $(NH_4)_2SO_4$, 0.01 g/L EDTA, 0.1 g/L $MgCl_2{\cdot}6H_2O$, 0.002 g/L $ZnSO_4{\cdot}7H_2O$, 0.001 g/L $CaCl_2{\cdot}2H_2O$, 0.005 g/L $FeSO_4{\cdot}7H_2O$, 0.0002 g/L $Na_2MoO_4{\cdot}2H_2O$, 0.0002 g/L $CuSO_4{\cdot}5H_2O$, 0.0004 g/L $CoCl_2{\cdot}6H_2O$, and 0.001 g/L $MnCl_2{\cdot}2H_2O$.

Glucose and sodium pyruvate stock solutions were prepared in Milli-Q water and sterilized by filtration through a 0.20-μm filter. Cumate and anhydrotetracycline (aTc) stock solutions were prepared in ethanol at concentrations of 100 mM and 1 mM, respectively, and sterilized by filtration through a 0.20-μm filter. At their maximum working concentrations of 100 μM cumate and 0.2 μM aTc, the corresponding final ethanol concentrations were 0.10% and 0.02% (v/v), respectively. Ampicillin and apramycin were used at final concentrations of 100 μg/mL and 50 μg/mL, respectively.

**Method S2. Data processing and mathematical modelling.**

Design of Experiments (DoE) data were analyzed using linear regression models. The relationship between the experimental response and the investigated factors was described using the following linear regression model (Eq. 1):

$$Y = \beta_0 + \sum_{i=1}^{n} \beta_i X_i + \sum_{i=1}^{n} \sum_{j=i+1}^{n} \beta_{ij} X_i X_j. \ (1)$$

Where $Y$ represents the predicted response, $X_i$ represents the considered factors, $\beta_0$ is the intercept, $\beta_i$ are the linear coefficient of the main effects (MEs), and $\beta_{ij}$ represents the coefficients of the two-factor interactions (2FIs). The quality of the regression equations was assessed according to the adjusted coefficient of determination (Adj $R^2$). Regression coefficients and their associated P values were used to assess the magnitude, direction, and statistical significance of the effects.

**Method S3. RNA secondary structure and complex prediction.**

The secondary structures of the toehold switch RNA, trigger RNA, and the toehold switch–trigger RNA complex were predicted using the NUPACK web application*(2)*. The minimum free energy (MFE) secondary structure and its corresponding free energy ($\Delta G$) were calculated using NUPACK. All simulations were performed with the RNA thermodynamic model at 37 °C. For single-strand predictions, the concentration of each RNA strand was set to 1 μM. To predict the toehold switch–trigger RNA complex, the concentrations of the toehold switch RNA and trigger RNA were each set to 1 μM, and the maximum complex size was set to 2.

**Method S4. Functional validation of the toehold switch system.**

GFP fluorescence was measured in *E. coli* BL21(DE3) harboring toehold switch constructs containing either a strong RBS or the weak RBS33 to assess the effect of RBS strength on system performance. The medium-strength $P_{J115}$ promoter was used to control expression of the toehold switch RNA, whereas the stronger $P_{J100}$ promoter was used to control expression of the trigger RNA. Strains were cultured in M9 medium containing 10 mM glucose and ampicillin.
